# The Abelson tyrosine kinase and the Nedd4-family ubiquitin ligase Suppressor of Deltex converge at the Notch PPxY motif to regulate endosomal trafficking and signaling

**DOI:** 10.1101/2020.10.20.347468

**Authors:** Nicelio Sanchez-Luege, Julio Miranda-Alban, Xiao Sun, Fernando M. Valbuena, Benjamin S. Glick, Ilaria Rebay

## Abstract

The conserved Notch signaling pathway coordinates diverse cellular processes during animal development. Unlike most cell surface receptors that use a cytoplasmic cascade to amplify and diversify signaling dynamics, Notch itself transduces external cues directly to the nucleus. How appropriate signaling dynamics and transcriptional responses are achieved with this pathway architecture remains unclear. Here, we report that the cytoplasmic tyrosine kinase Abelson (Abl) fine-tunes Notch signaling by regulating Notch endocytic trafficking. We show that Abl can directly phosphorylate a PPxY motif important for Nedd4-family ubiquitin-ligase-mediated transfer of Notch into degradative endosomal compartments. Consistent with this, loss of Abl or inhibition of its kinase activity results in aberrant endosomal accumulation of Notch, while mutation of the PPxY tyrosine renders Notch insensitive to such regulation. Phenotypic and genetic interaction studies in the wing, together with parallel assays in cultured cells, show that loss or gain of Abl activity can respectively increase or decrease Notch output. We propose that the Notch PPxY motif operates as a molecular hub that integrates multiple post-translational modifications to regulate Notch trafficking and fine-tune signaling output.

## Introduction

Development depends on precisely orchestrated patterns of cell survival, proliferation, fate specification, and morphogenesis. While the core molecular signaling events that trigger particular cell fate transitions have been intensively studied, the mechanisms that fine-tune signaling activity in different contexts are less well understood. The Notch pathway provides an ideal system to investigate the mechanisms that regulate the initiation, strength and duration of signaling during development (Hori et al., 2013). Animals broadly express the Notch receptor during development and spatiotemporal Notch activation drives a plethora of cellular transitions, ranging from neurogenesis to programmed cell death. Notch activation must be precisely regulated, with both too much or too little signaling producing developmental defects depending on the spatio-temporal context (Aster et al., 2017; Zacharioudaki and Bray, 2014). In this study, we investigate how the Drosophila Abelson (Abl) cytoplasmic tyrosine kinase regulates Notch signaling by affecting its trafficking through endosomal compartments.

Notch is an integral membrane protein with a ligand-binding extracellular domain and a signal transducing intracellular domain (Wharton et al., 1985). Proteolytic cleavage of the Notch intracellular domain (NICD) marks the fundamental molecular signaling event (Hori et al., 2013; Lieber et al., 1993; Rebay et al., 1993). Free NICD enters the nucleus and forms a transcriptional complex that drives target gene expression (Bailey and Posakony, 1995; Wilson and Kovall, 2006; Wu et al., 2000). In ligand-dependent activation, Notch receptor binds Delta/Serrate/Lag2-family ligands exposed on neighboring cells (Gordon et al., 2008). Ligand recognition triggers cleavage of the Notch extracellular domain (NECD) by the proteinase ADAM, a process called S2 cleavage (Brou et al., 2000). Following S2 cleavage, another cleavage site is exposed which triggers cleavage and release of NICD by γ-secretase and presenilin, deemed S3 cleavage (Strooper et al., 1999). These events occur within endocytic vesicles following ligand-mediated endocytosis (Yamamoto et al., 2010).

Pulse chase experiments have shown that cells continuously traffic Notch from the cell surface into endocytic compartments (Maitra et al., 2006). Endocytosed Notch can either be recycled back to the surface, or degraded within the endocytic pathway (Yamamoto et al., 2010). When degraded, Notch is sorted through early endosomes, multivesicular bodies, and the lysosome (Vaccari et al., 2008; Yamamoto et al., 2010). Mutating ESCRT complex factors, which regulate internalization of proteins into the multivesicular body (Henne et al., 2011), causes Notch accumulation and activation in the absence of detectable ligand expression (Vaccari et al., 2008). Therefore, Notch trafficking is actively regulated both to prevent ligand-independent activation and to fine-tune ligand-dependent signaling.

Notch trafficking is regulated by ubiquitination (Hori et al., 2013; Shimizu et al., 2014), a post-translational regulatory mechanism that modulates endocytic trafficking and degradation of many receptors (Hershko and Ciechanover, 1998). Important insights have come from studying two genetic modifiers of Notch that both encode ubiquitin ligases: Deltex (Dx) and Suppressor of dx (Su(dx)). Genetic and molecular studies implicate Dx as a positive regulator and Su(dx) as a negative regulator of Notch signaling (Fostier et al., 1998; Fuwa et al., 2006; Shimizu et al., 2014). *dx* was originally identified as a spontaneous mutant that both recapitulates and strongly enhances Notch loss-of-function phenotypes (Xu and Artavanis-Tsakonas, 1990). Dx loss reduces Notch endocytosis, and its overexpression can promote endocytosis and ligand-independent activation (Hori et al., 2011; Yamada et al., 2011; Shimizu et al., 2014). Molecularly, Dx operates as a RING family ubiquitin ligase that ubiquitinates the NICD and directs endocytosed Notch to the lysosome (Hori et al., 2004; Wilkin et al., 2008). *Su(dx)* was discovered as a spontaneous suppressor of the Notch-like deltex phenotypes (Fostier et al., 1998) and encodes a member of the Nedd4-family HECT ubiquitin ligases (Cornell et al., 1999). Mechanistically, Su(dx) has two distinct and molecularly separable activities in Notch regulation. First, Su(dx) promotes endocytosis of Notch into a sterol-rich endocytic network via a ubiquitination-independent mechanism (Shimizu et al., 2014). Second, via an interaction between the WW domains of Su(dx) and a proline-rich motif in the NICD, Su(dx) ubiquitinates Notch to promote internalization into the multivesicular body (MVB) (Sakata et al., 2004). This topologically restricts cleaved NICD from accessing the nucleus and thus represses signaling.

Previous studies of Abl function during photoreceptor morphogenesis serendipitously identified it as a candidate regulator of Notch trafficking and signaling: loss of Abl resulted in a failure to clear endocytosed Notch and an increase in Notch target expression, while Notch pathway mutations dominantly suppressed *abl* mutant phenotypes (Xiong et al., 2013). In this study we use established *in vivo* and cultured cell models to explore the molecular and genetic mechanisms of Abl-mediated regulation of Notch trafficking and signaling. We first demonstrate that Abl promotes Notch clearance from endosomes in a kinase-dependent way and that Abl can phosphorylate the PPxY tyrosine in the NICD. We further find that the PPxY tyrosine is required both for regulation of Notch trafficking by Su(dx) and for the negative regulation of Notch signaling by Abl and Su(dx). Altogether, these data support a mechanistic model in which Abl-directed phosphorylation of the Notch PPxY domain promotes Su(dx)-mediated internalization of Notch into the multivesicular body (MVB) to limit signaling output. We suggest that adjusting the balance of endocytosed Notch between different endosomal compartments may be broadly applied as a molecular strategy to fine-tune Notch signaling.

## Results

### Abl prevents endosomal Notch accumulation in a kinase-dependent way

As previously reported (Xiong et al., 2013), retinal cells lacking Abl fail to clear endocytosed Notch, resulting in an increased accumulation of cytoplasmic Notch (Figure 1A,B). The same is observed in *abl* mutant wings (Figure 1C,D), suggesting that Abl regulates Notch trafficking more broadly during development (Xiong et al., 2013). If Abl normally promotes clearance of endocytosed Notch to limit signaling (Xiong et al., 2013), then Abl loss should cause Notch to accumulate in signaling competent compartments defined by previous studies (Vaccari and Bilder, 2005; Vaccari et al., 2008). To test this we stained *abl* clones for well-characterized markers of early, mid and late endocytic compartments: Eps15, a marker of upstream endocytic cargo vesicles (Tebar et al., 1996), Hrs, a marker of early endosomes (Raiborg et al., 2002), and Rab7, a late endosomal marker (Bucci et al., 2000). Notch puncta colocalized with Hrs in *abl* clones (Figure 2B), but did not colocalize with Eps15 (Figure 2A) and only sparingly colocalized with Rab7 (Figure 2C). The overall intensity and pattern of all three markers was indistinguishable between *abl* mutant and wild type tissue (Figure S2A-C), arguing that the excessive accumulation of Notch into early endosomes under Abl loss is unlikely the result of a grossly disrupted endocytosis.

**Figure 1.**
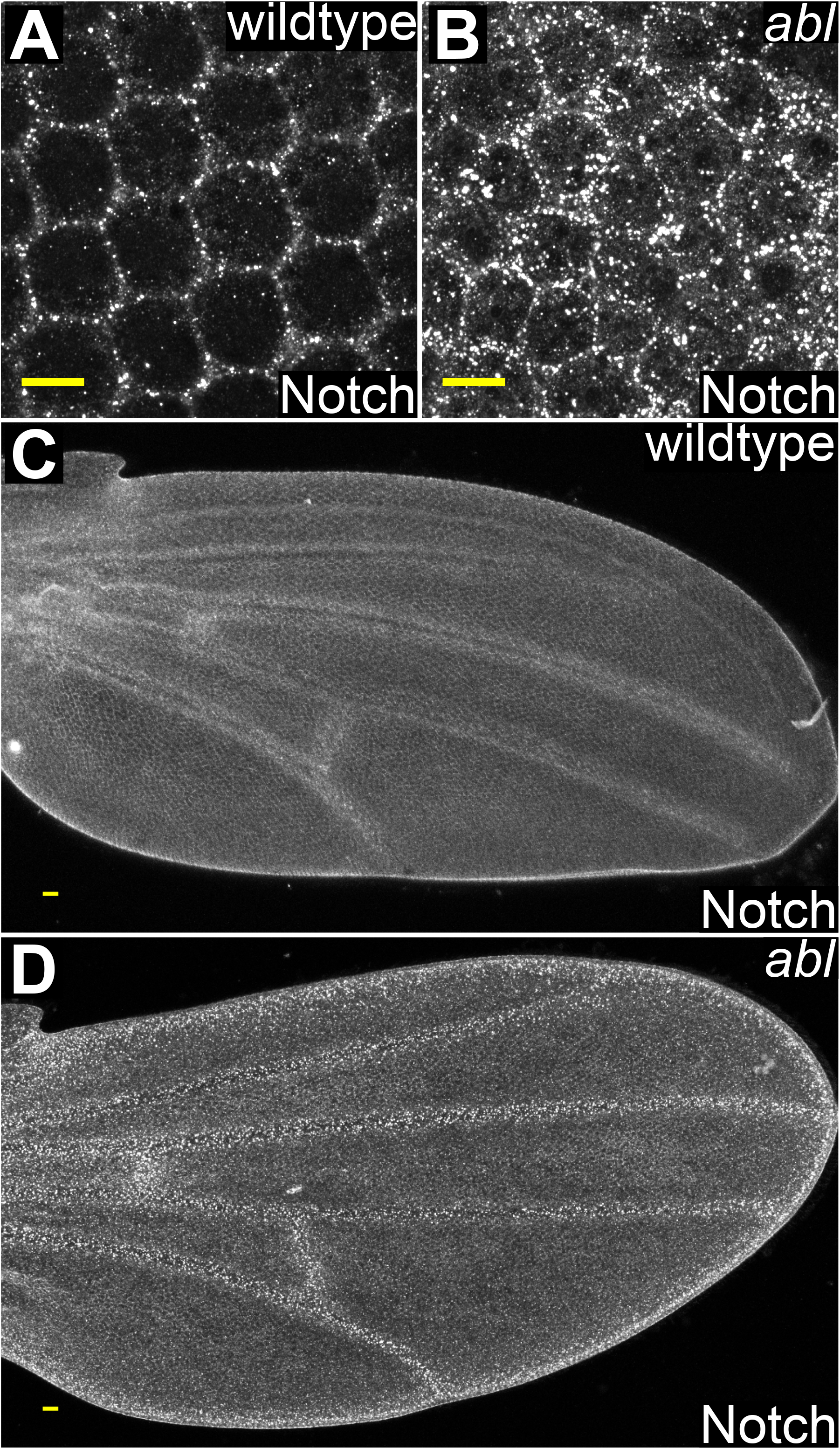
Abl prevents Notch accumulation. (A, B) Panels show maximal confocal projections of 48h APF retinas. Notch accumulates into prominent puncta in *abl* mutant retinas. Scale bar = 10 μm. (C-D) Panels show maximal confocal projections of 32h APF wings. Scale bar = 10 μm (C) Notch is diffusely expressed with enrichment in the flaking inter-vein regions. (D) Notch forms distinct large puncta in *abl* mutants.

**Figure 2.**
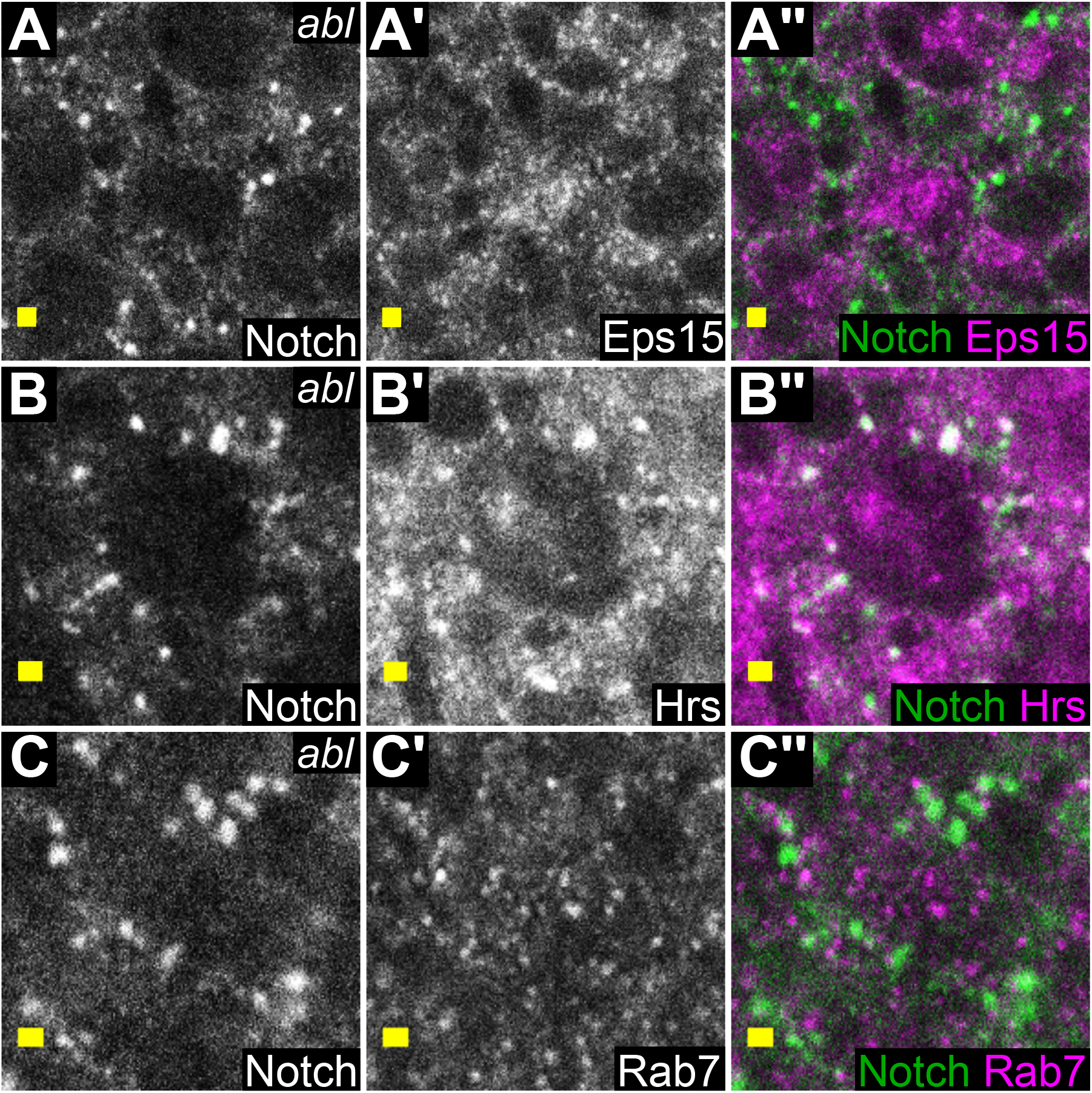
Notch accumulates in Hrs+ compartments in *abl* mutants. (A-C) Panels show single confocal sections of *abl* mutant ommatidia at 48h APF. Scale bar = 1μm. (A-A’’) Notch puncta do not overlap with Eps15 puncta. (B-B”) Notch puncta extensively overlap with Hrs puncta (C-C’’) Notch puncta have limited overlap with Rab7 puncta.

To address whether the mechanism by which Abl limits Notch accumulation in early endosomes is kinase-dependent, we transfected Notch into S2 cells, which express Abl endogenously (Zhang et al., 2010) but do not express Notch (Fehon et al., 1990), and treated the cells with 200 μM imatinib mesylate, an Abl kinase activity inhibitor (Hunter, 2007). In untreated cells, Notch localized to the cell membrane, diffusely within the cytoplasm, and in distinct puncta (Figure 3A). Under imatinib mesylate treatment, Notch concentrated into large cytoplasmic puncta (Figure 3B), similar to the effect of Abl loss *in vivo.* These structures were Hrs+ (Figure S3A), again consistent with the pattern of Notch accumulation in *abl* mutant tissue.

**Figure 3.**
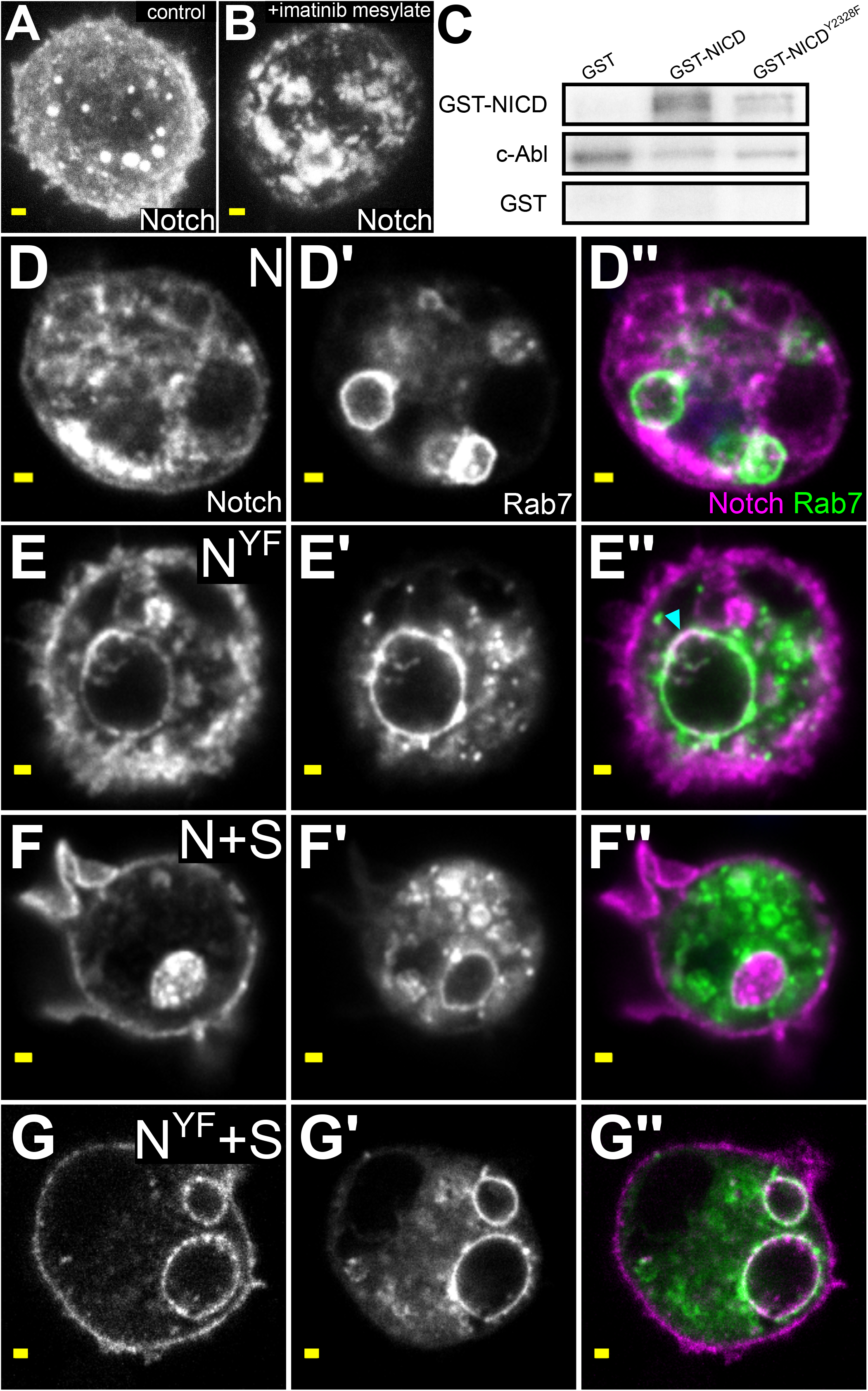
Abl regulates Notch trafficking via the PPxY motif. (A-B) Panels show maximal confocal projections of representative S2 cells expressing Notch. Scale bar = 1 μm. (A) An untreated cell with Notch at the cell membrane, diffusely in the cytoplasm, and concentrated into small puncta. (B) A cell treated with 200 μM imatinib mesylate for 24 hours. Notch localizes into large cytoplasmic puncta. (C) Recombinant murine Abl phosphorylates the NICD with some activity targeted to Y2328. In lane 1, Abl phosphorylates itself but not GST alone. In lane 2, Abl phosphorylates itself and GST-NICD. In lane 3, Abl phosphorylates itself with slightly higher intensity as lane 2. However, Abl phosphorylates GST-NICD^Y2328F^ with less intensity than in lane 2, indicating that at least some Abl activity targets Y2328. Abl auto-phosphorylation appears more intense than in lane 2. (D-G) All panels show single-slice confocal images of representative S2 cells expressing Notch and the late-endosomal marker Rab7-GFP.. Scale bar = 1μm. (D-D’’) Notch does not colocalize with late endosomes when expressed alone. (E-E’’) Notch^Y2328F^ localizes to the limiting membrane of late endosomes, cyan triangle. (F-F’’) Notch is internalized into late endosomes when coexpressed with Su(dx). (G-G’’) Notch^Y2328F^ localizes to the limiting membrane of late endosomes when coexpressed with Su(dx).

### Abl phosphorylates the Notch PPxY motif *in vitro*

Does Abl directly phosphorylate Notch to prevent its accumulation into early endosomes? The Drosophila NICD contains 15 tyrosine residues (Wharton et al., 1985). One residue, Y2328, is within the PPxY motif recognized by Nedd4-family ubiquitin ligases (Jennings et al., 2007). Given that Su(dx) and Nedd4, the Drosophila Nedd4-family ubiquitin ligases, also regulate Notch trafficking and signaling (Wilkin et al., 2004), we wondered whether Abl can phosphorylate Notch at the PPxY tyrosine and if so, whether this might impact Su(dx)-mediated regulation. Using *in vitro* kinase assays, we found that Abl phosphorylated GST-NICD, but not GST alone (Figure 3C). Visibly less phosphorylation was detected when GST-NICD^Y2328F^ was used as the substrate (Figure 3C, S3B), indicating that Y2328 can be a direct target of Abl activity.

### The Notch PPxY tyrosine is required for endosomal internalization by Su(dx)

The ability of Abl to phosphorylate Notch at the PPxY domain offered a new mechanistic model for exploring how Abl kinase activity regulates Notch. To start we reasoned that if the endocytic accumulation of Notch that occurs in *abl* mutants *in vivo* results from loss of regulation at this tyrosine residue, then the subcellular localization of wild type and Y2328F mutant Notch should be different. Comparison of wild type and Y2328F mutant Notch in transiently transfected S2 cells revealed both similarities and differences in subcellular localization. Both showed comparable cytoplasmic and plasma membrane-associated localization (Figure 3D,E). However, only Notch localized to the outside of distinct vacuolar structures that co-transfection with Rab7-GFP identified as late endosomes (Figure 3E-E”, cyan triangle).

Prior studies have shown that Su(dx) promotes internalization of Notch into the late endosome lumen (Shimizu et al., 2014; Wilkin et al., 2004). Therefore if Notch^Y2328F^ mislocalizes to the limiting membrane of late endosomes as suggested by its colocalization with Rab7-GFP, then it should also be resistant to Su(dx). To test this we co-transfected Su(dx) and again compared the subcellular localization of Notch versus Notch^Y2328F^. Whereas co transfection of Su(dx) caused the expected internalization of wild type Notch into the lumen of Rab7-GFP marked compartments (Figure 3F-F”), Notch^Y2328F^ remained at the limiting membrane (Figure 3G-G”). Therefore, the Y2328F mutation prevents Su(dx)-mediated internalization into late endosomes. By extension, phosphorylation of Y2328, perhaps mediated by Abl, may facilitate Notch internalization into late endosomes.

### The Notch PPxY tyrosine is required for regulation of Notch signaling by Abl and Su(dx)

Transfer of Notch to internal endosomal compartments, as promoted by Su(dx), limits Notch signaling by topologically sequestering cleaved NICD within the endosome (Sakata et al., 2004; Wilkin et al., 2004). Therefore the mislocalization of Notch^Y2328F^ to the limiting membrane of late endosomes and its insensitivity to Su(dx) predict it should have intrinsically elevated signaling activity. Using a ligand-independent assay of Notch pathway transcriptional output in transiently transfected cultured S2 cells (Zacharioudaki and Bray, 2014), we found that Notch^Y2328F^ exhibited approximately twofold greater activity than wildtype Notch (Figure 4A, p = 2.98×10^-4^). This shows that the PPxY tyrosine normally limits Notch signaling output. Because PPxY also provides a recognition motif for Su(dx), and because like Abl, Su(dx) exhibits a genetically antagonistic relationship with Notch (Fostier et al., 1998), the increased activity of Notch^Y2328F^ may result from compromised interaction with Abl, with Su(dx), or with both.

**Figure 4.**
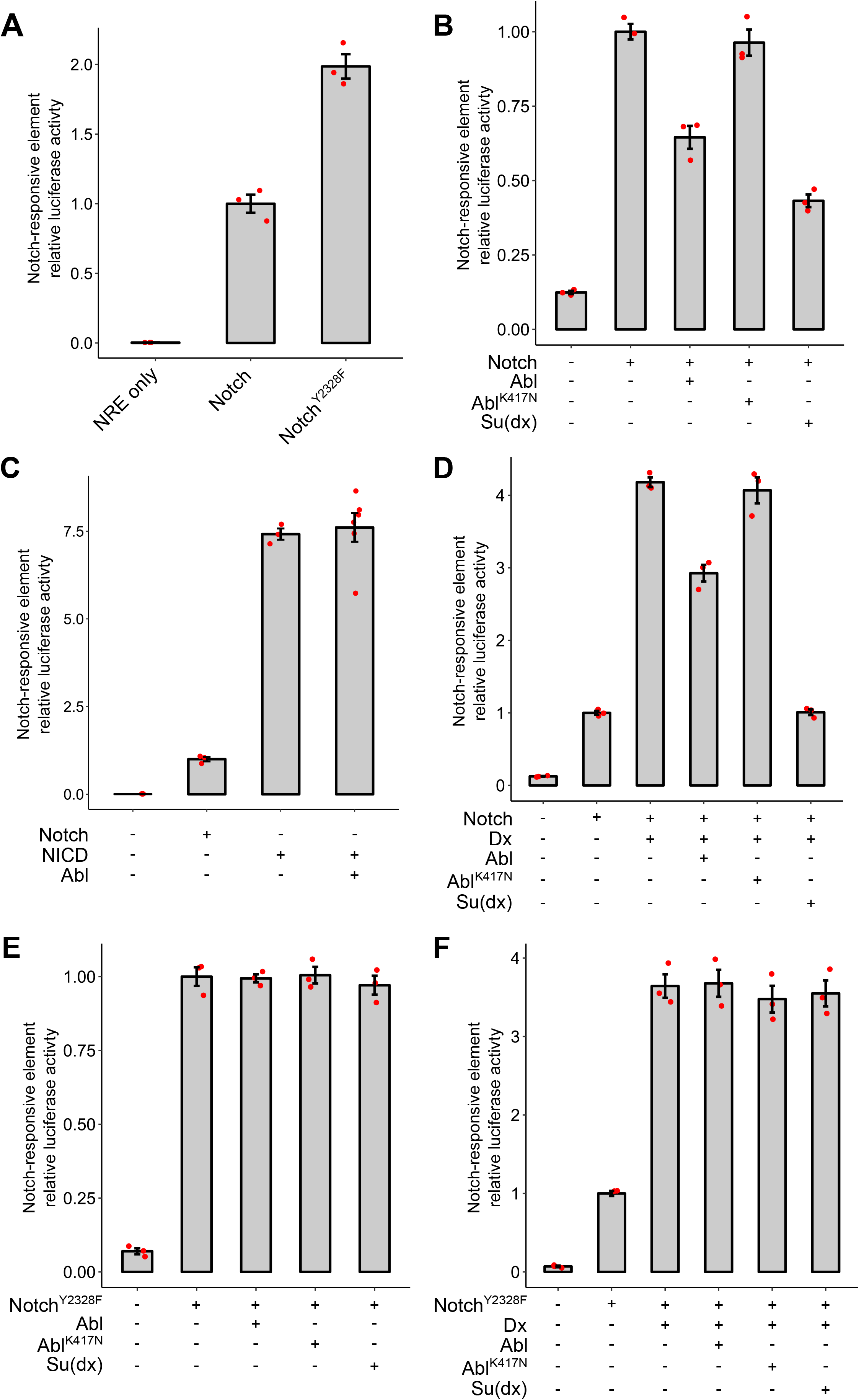
The Notch PPxY tyrosine is required for regulation of Notch signaling by Abl and Su(dx) All panels show quantification of S2 cell transcription assays. Each red dot represents a biological replica. (A) Notch^Y2328F^ exhibits ~2-fold greater signaling activity compared to wildtype Notch. (p = 2.98×10^-4^) (B) Notch activity is reduced by expression of Abl (p = 0.0026) or Su(dx) (p = 9.67 x 10^-5^), but not the kinase-deficient mutant Abl^K417N^ (p = 0.51). (C) NICD activity is unaffected by Abl expression. (p = 0.67) (D) Dx-enhanced Notch activity is reduced by Abl (p = 0.0018) or Su(dx) (p = 1.52 x 10^-5^), but not Abl^K417N^ (p = 0.60). (E) Notch^Y2328F^ activity is unaffected by expression of Abl (p = 0.88), Su(dx) (p = 0.55), or Abl^K417N^ (p = 0.91). (F) Dx-enhanced Notch^Y2328F^ activity is unaffected by expression of Abl (p = 0.88) or Su(dx) (p = 0.70).

The model that Abl kinase activity antagonizes Notch signaling by affecting its endosomal trafficking makes two predictions: 1) that a kinase-deficient Abl mutant should fail to regulate Notch signaling, and 2) that Abl should fail to regulate the signaling activity of free NICD. To test these predictions, we asked how co-transfection of Abl altered Notch signaling output in the S2 cell reporter assay. Abl overexpression significantly decreased Notch activity (Figure 4B, p = 0.0026), consistent with previously reported genetic antagonism (Xiong et al., 2013). In contrast, co-transfection of Abl^K417N^, a kinase-deficient mutant (Henkemeyer et al., 1990), had no significant effect on Notch activity (Figure 4B, p = 0.51), suggesting that Abl indeed regulates Notch signaling in a kinase-dependent way. This regulation is directed at fulllength Notch, since the signaling activity produced by expression of free NICD was unaffected by Abl expression (Figure 4C, p = 0.67). These results suggest that Abl kinase activity regulates Notch signaling prior to release of NICD.

### Abl antagonizes Dx-mediated Notch activation

A previous study found that Su(dx) buffers Notch signaling against excessive activation by Dx (Shimizu et al., 2014, Figure 4D, p = 1.53 x 10^-5^). Therefore, if Abl promotes this Su(dx)-Notch interaction, then Abl should also render Notch less susceptible to activation by Dx. To test this, we co-transfected Dx and measured the effect on Notch signaling output. Dx expression increased Notch activity approximately 4-fold compared to Notch alone (Figure 4D, p = 7.82 x 10^-5^). Transfection of Abl in this Dx-enhanced background significantly reduced reporter activity (p = 0.0018), while Abl^K417N^ had no significant effect (p = 0.60). Therefore, Abl can limit Dx-mediated induction of Notch signaling in a kinase-dependent way.

The model that Abl specifically targets Y2328 predicts that Notch^Y2328F^ should be resistant to Abl activity. Emphasizing the importance of this tyrosine to Abl-mediated Notch regulation, the increased signaling output from Notch^Y2328F^ was not significantly affected by cotransfection of either Abl or Abl^K417N^ (Figures 4E, p = 0.88 and 0.91 respectively). Notch^Y2328F^ was also insensitive to Su(dx) (Figure 4E, p = 0.55), whereas activation of wildtype Notch was attenuated by Su(dx) expression (Figure 4B, p = 9.67×10^-5^). Finally, while cotransfection of Dx increased Notch^Y2328F^ about 3.5 fold (p = 0.0022), co-transfection of neither Abl nor Su(dx) reduced signaling output (Figure 4F, p = 0.88 and 0.70 respectively). Together these results suggest that both Abl and Su(dx) target Y2328 to regulate signaling, and that this regulation can limit Notch signaling against excessive Dx-driven activation.

### Abl antagonizes Notch activation *in vivo*

We next asked whether Abl regulates *in vivo* Notch signaling in a way consistent with the results of cell culture transcriptional assays (Figure 4B). We focused on wing vein development, the context in which Su(dx)-mediated antagonism of Notch function was initially described (Cornell et al., 1999; Fostier et al., 1998). We found that primordial veins, marked by Delta, were truncated in *abl* mutants during pupal wing development (Figure 5A, B, cyan triangle), a phenotype consistent with overactivation of Notch (Celis et al., 1997). Furthermore, overexpression of Abl using the *apterous* promoter caused vein thickening compared to *white^1118^* conrols (Figure 5C, D, blue triangle), consistent with a decrease in Notch signal (Zacharioudaki and Bray, 2014). Overexpression of Abl^K417N^ did not cause vein thickening (Figure 5E) and produced wings indistinguishable from controls, suggesting that Abl overexpression decreases Notch signaling in a kinase-dependent way *in vivo*, consistent with the cell culture results.

**Figure 5.**
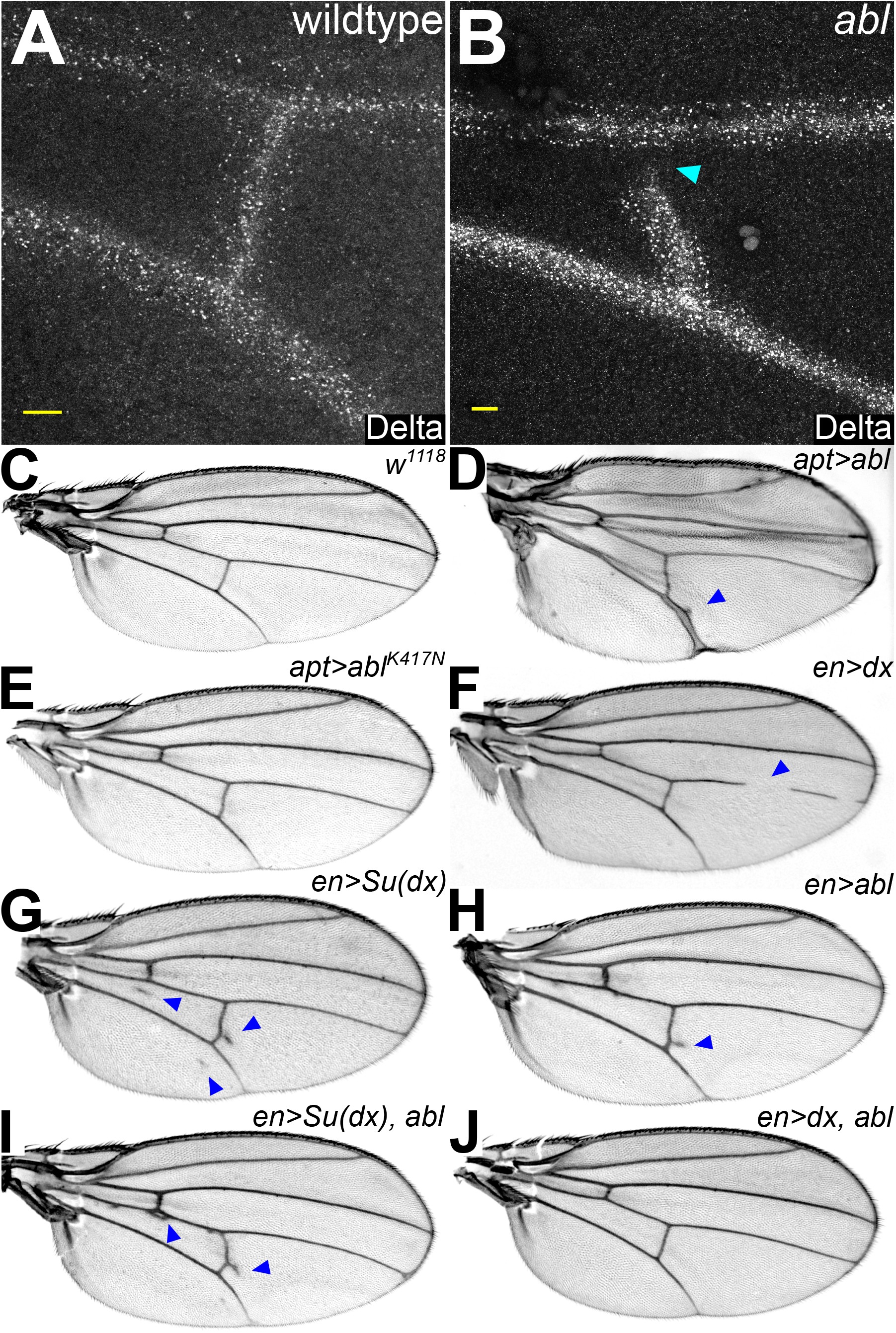
Abl regulates Notch signaling and genetically antagonizes Dx *in vivo*. (A-B) Panels show maximal confocal projections of primordial posterior crossveins from 32h APF wings stained for Delta. Scale bar = 10 μm. (A) A wild type wing with a normally intact posterior cross vein. (B) An *abl* mutant wing with a truncated cross vein marked by a cyan triangle. (C-I) Light microscope images of representative adult wings. (C) *white^1118^* shows the wildtype vein phenotype (D) *apt>abl* with longitudinal vein thickening, blue triangle (E) *apt>abl^K417N^*, indistinguishable from wild type (F) *en>dx* with longitudinal vein truncation, blue triangle. (G) *en>Su(dx)* with ectopic vein tissue, blue triangles (H) *en>abl* with ectopic vein tissue, blue triangles (I) *en>Su(dx), abl* with enhanced ectopic vein tissue, blue triangles (J) *en>dx, abl* with mutual suppression of ectopic vein tissue and vein truncation.

To test whether Abl exhibits the expected genetic synergy with Su(dx) and antagonism with Dx, we overexpressed them along with Abl in the posterior compartment of the wing during pupation using the *engrailed* promoter for spatial control and the temperature-sensitive Gal80 repressor for temporal control (McGuire et al., 2004; see methods for details). Overexpression of Dx resulted in vein truncation at 63% (n = 22) penetrance (Figure 5F), consistent with its known role as a Notch activator (Matsuno et al., 1995). In contrast, overexpression of Su(dx) resulted in ectopic vein formation at 67% (n = 24) penetrance (Figure 5G). Overexpression of Abl also caused ectopic vein formation at 54% (n = 55) penetrance (Figure 5H). Therefore, Abl and Su(dx) both exhibit effects consistent with repression of Notch signaling. Simultaneous overexpression of Abl and Su(dx) resulted in ectopic vein formation at 86% (n = 83) penetrance, an enhancement over Abl overexpression alone (p = 7.95 x 10^-5^) (Figure 5I). In contrast, simultaneous overexpression of Abl and Dx results in vein truncation at 21% (n = 24) penetrance and ectopic vein formation at 5% penetrance (Figure 5J). Therefore, Abl and Dx mutually suppress their respective effects on pupal wing development (p = 1.10 x 10^-5^ and 0.0063 respectively), suggesting that Abl can antagonize excessive Notch activation driven by Dx *in vivo*.

### Loss of Abl diverts Su(dx) to its ubiquitin-ligase-independent function of promoting Notch endocytosis

Another prediction of the model that Abl phosphorylates Y2328 to promote Su(dx) activity is that Abl loss should impair the ubiquitin-ligase dependent activity of Su(dx) to internalize Notch into the MVB, but the ubiquitin-ligase independent activity of Su(dx) to promote Notch endocytosis should remain intact. In the absence of Abl, overexpression of Su(dx) would increase Notch endocytosis without the balance of increased internalization and degradation. Therefore, an even greater aberrant accumulation of cytoplasmic Notch should occur in an *abl* mutant background. To test this, we overexpressed Su(dx) in the background of *abl* pupal eye clones using the pan-eye GMR-Gal4 driver. Su(dx) overexpression visually increased Notch accumulation in *abl* clones (Figure 6A-B). Quantification showed that Notch puncta were 2.7-fold larger in *abl* clones under Su(dx) overexpression than in *abl* clones without Su(dx) overexpression (p = 9.07 x 10^-6^), while Su(dx) overexpression did not change Notch punctual size or accumulation in otherwise wildtype tissue (p = 0.77) (Figure 6C). This is consistent with the model that Abl promotes effective Su(dx)-mediated internalization and degradation of Notch within endosomal vesicles. Cells lacking Abl are highly sensitized to Notch endocytosis, suggesting a role for Abl in balancing the flux of Notch through endocytic compartments.

**Figure 6.**
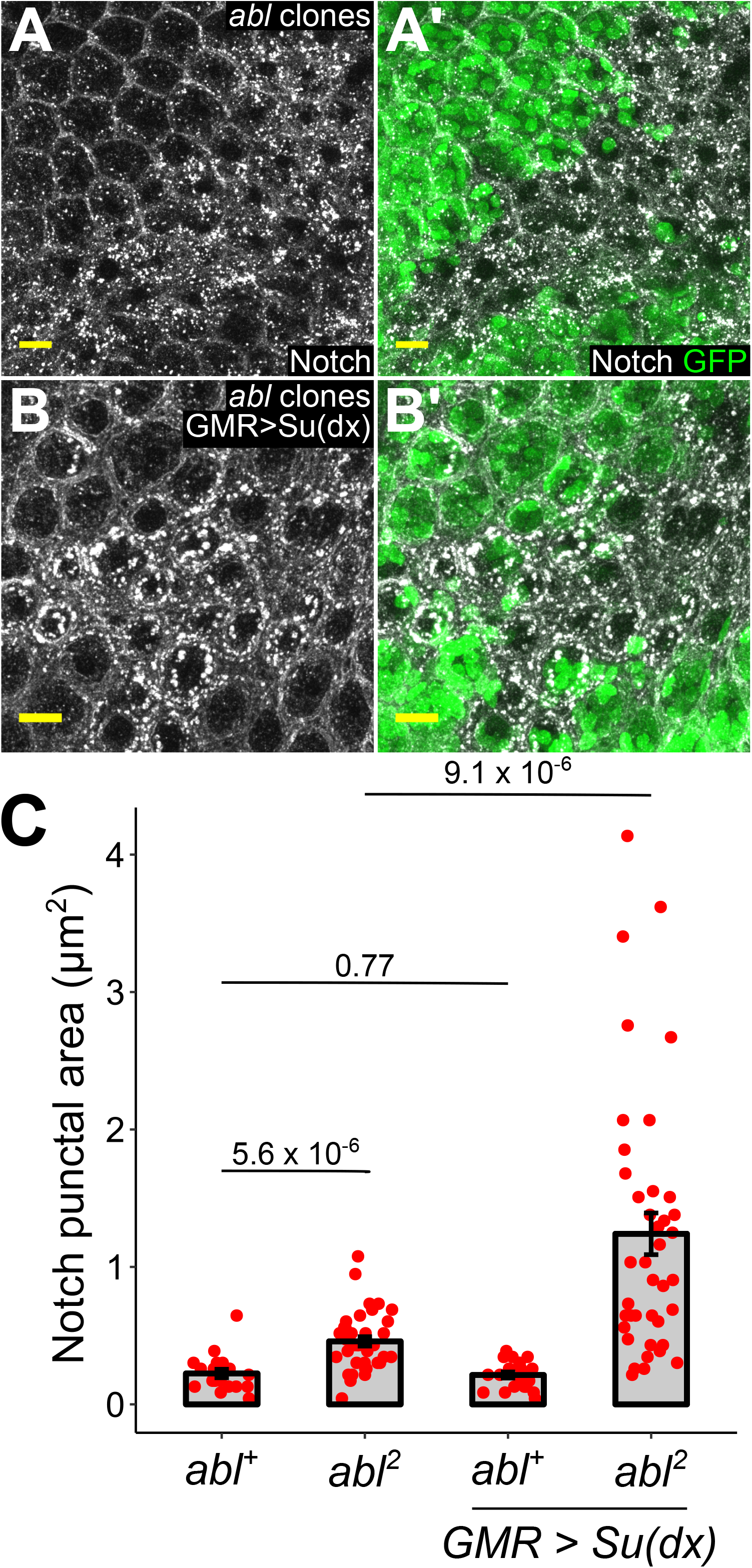
Loss of Abl diverts Su(dx) to its ubiquitin-ligase-independent function of promoting Notch endocytosis. (A-B) Panels show maximal confocal projections from 48h APF retinas containing *abl* mutant clones. Clones are marked by the absence of GFP. Scale bar = 10 μm. (A-A’) Notch accumulates into visibly larger puncta in *abl* clones. (B-B’) Overexpression of Su(dx) enhances the size of Notch puncta in *abl* clones but has no effect on punctal size in wildtype tissue. (C) Quantification of the findings in A and B, calculated as the maximal cross-sectional area of Notch puncta within each genetic category. Comparison t-test p-values are indicated within the panel.

## Discussion

In this study we establish the Abl non-receptor tyrosine kinase as a novel regulator of Notch trafficking and signaling. We show that Abl promotes clearance of Notch from endosomal compartments in a kinase dependent manner, and that it can phosphorylate the Notch PPxY motif, an important site for Nedd4 ubiquitin ligase-mediated regulation. Functional assays in cultured cells and in fly tissues suggest that the combined action of Abl and Su(dx) provides a molecular buffering mechanism that directs Notch transit through the endocytic pathway to prevent excessive Notch activation. More broadly we propose that the PPxY motif serves as a versatile molecular hub that integrates tyrosine phosphorylation and WW domain based regulatory inputs to optimize Notch signaling output.

Connections between receptor internalization, endocytic trafficking, and Notch signaling were first noted over 25 years ago (Fehon et al., 1990, 1991; Henderson et al., 1994; Kooh et al., 1993; Parks et al., 2000; Seugnet et al., 1997). More recent studies have demonstrated the importance of Nedd4 ubiquitin ligases in orchestrating Notch endocytic dynamics (Conner, 2016; Sakata et al., 2004; Shimizu et al., 2014; Wilkin et al., 2004; Yamada et al., 2011). In terms of general interaction mechanism, Nedd4 ligases contain WW protein domains that recognize and bind PPxY motifs in their substrates (Otte et al, 2002; Ingham et al., 2004). For example, nuclear magnetic resonance revealed the structure of a Su(dx) WW domain bound to the Drosophila NICD PPxY motif (Jennings et al., 2007). Biochemical analyses of either Drosophila Nedd4-Notch complexes or the analogous human WWP2-NOTCH3 interaction have shown that mutation of the PPxY tyrosine reduces Nedd4 binding and ubiquitination of Notch/NOTCH3, although the downstream impact on signaling was not explored (Jung et al., 2014; Sakata et al., 2004). Our study has now revealed a functional consequence of disrupting the PPxY motif, namely that Notch^Y2328F^ is resistant to both Abl-mediated regulation and Su(dx)-driven endosomal internalization and therefore has stronger signaling output.

Although structural changes could underlie the insensitivity of Notch^Y2328F^ to both Abl and Su(dx), prior work demonstrating the general importance of PPxY tyrosines in mediating interaction with WW domain proteins (Kasanov et al., 2001; Liu et al., 2016; Otte et al., 2003) leads us to propose that the phosphorylation state of the Notch PPxY motif regulates its interaction with Su(dx). These *in vitro* screens have shown that WW domains can bind both unphosphorylated and tyrosine phosphorylated PPxY motifs, with phosphorylation preference depending on the specific WW domain and the experimental conditions (Liu et al., 2016; Otte et al., 2003). Given the lack of general rule for how PPxY phosphorylation state impacts the binding preference of Nedd4-family ubiquitin ligases, the results of our study can be interpreted with two mutually non-exclusive mechanistic models.

One model is a simple serial interaction in which Abl first phosphorylates the NICD PPxY tyrosine, which promotes Su(dx) recruitment and increases the probability of ubiquitination and degradation of Notch in signaling-incompetent compartments. An analogous linear model, but with the opposite relationship in which PPxY phosphorylation prevents Nedd4-family ligase binding, was previously proposed to regulate the activity of the transcription factor c-Jun in T-cells (Gao et al., 2006). c-Abl was implicated as the responsible kinase in that study, affirming its general ability to recognize and phosphorylate PPxY motifs.

The second model is that the PPxY provides a molecular hub whose phosphorylation status can be independently read and acted upon by many different classes of proteins including tyrosine kinases, tyrosine phosphatases, SH2 or PTP phosphotyrosine-binding domaincontaining proteins and WW domain-containing proteins. Some of these interactions would be serially coupled, while others would occur in parallel. With regard to Su(dx), interaction with PPxY versus PPxYp might direct Su(dx) to apply different ubiquitination patterns on NICD, resulting in different trafficking routes. For example, mono-versus polyubiquitination by the Nedd4-family yeast ortholog Rsp5 functions as a nitrogen-sensing switch that trafficks the transmembrane substrate to the plasma membrane or the vacuole, respectively (Helliwell et al., 2001). A similar switch behavior in which monoubiquitinated Notch is recycled to the plasma membrane could explain why Su(dx) overexpression enhances Notch accumulation under Abl loss (Figure 6). The potential for the phosphorylation state of the PPxY motif to modulate a complex network of protein-protein interactions thus offers a rich regulatory repertoire to influence Notch trafficking and fine-tune signaling activity.

How might this regulation apply to Notch signaling in vivo? One possibility is that the phosphorylation state of the NICD PPxY is spatiotemporally patterned during development to maintain appropriate Notch signaling levels. Consistent with this idea, Abl expression is enriched in the inter-vein regions flanking the primordial veins during pupal wing development (Bennett and Hoffmann, 1992). Su(dx) expression is similarly enriched in this flanking inter-vein region (Cornell et al., 1999). Together these patterns suggest that tight regulation of endocytic trafficking may be critical to attenuating Notch signaling in cells that are exposed to high concentration of Delta ligand, which is expressed in the primordial veins. This regulation is unlikely to be at saturation, since overexpressing Abl during wing development produces vein thickening consistent with Notch signal attenuation (Figure 5D).

Spatiotemporal patterning of NICD phosphorylation raises the question of whether phosphorylation confers an effect beyond trafficking of the full-length receptor. Our finding that NICD^Y2328F^ has no difference in signaling output relative to NICD suggests that biochemical regulation of full-length Notch trafficking is decoupled from downstream transcriptional activity. This is consistent with the conventional view that individual NICD molecules are interchangeable with respect to their transcriptional activity within the nucleus. Alternatively, a molecular memory of pre-cleavage events could add yet another layer of complexity to the spatiotemporal regulation of Notch signaling. Different trafficking events, biochemically marked by post-translational modification of the NICD, could activate distinct target genes or produce different transcriptional dynamics. A recent study of transcriptional dynamics downstream of Notch signaling proposed that NICD levels control the duration of transcriptional bursts at Notch target genes (Falo-Sanjuan et al., 2019). Whether tyrosine phosphorylation or other modifications of NICD alter transcriptional dynamics at individual Notch-responsive enhancers remains an open question.

## Materials and Methods

### Fly genetics

All strains were from the Bloomington stock center (BSC) or derived from BSC lines unless otherwise noted. All crosses were performed at 25C unless otherwise noted. To generate *abl* retinal clones, *abl^2^, FRT80B/TM6, Tb* males were crossed to *ey-flp; ubi-GFP, FRT80B* females. To generate *abl^2^* retinal clones with marked Rab7, *abl^2^, FRT80B, rab7-myc-YFP/TM6, Tb* males were crossed to *ey-Flp; ubi-GFP, FRT80B* females. To generate whole mutant *abl* retinas or wings, *abl^1^/TM6, Tb* males were crossed to *abl^2^, FRT80B/TM6, Tb* females. To generate *abl* retinal clones that overexpress Su(dx), *GMR>Su(dx)/Cyo, actin-GFP; abl^2^, FRT80B/TM6, Tb* males were crossed to *ey-Flp; ubi-GFP, FRT80B* females. To evaluate the effect of Abl overexpression on wing development *UAS-Abl-GFP* and *UAS-AblK417N-GFP* (O’Donnell and Bashaw, 2013) females were crossed to *apt-Gal4* males. To evaluate genetic interactions between Abl, Dx and Su(dx) in the wing, *en-Gal4/Cyo, actin-GFP; tubGal80^ts^/TM6, Tb* females were crossed to males of the following lines: *UAS-Abl-GFP, UAS-Su(dx), UAS-Flag-Dx, UAS-Su(dx); UAS-Abl-GFP* and *UAS-Flag-Dx; UAS-Abl-GFP*. Crosses were established at 18C and individual animals were transferred to 25C at 0h APF.

### Immunostaining and antibodies

48 hours APF retinas were dissected in PBS and fixed for 10 minutes in 4% paraformaldehyde in PBS with 0.1% Triton X-100 (PFA+PBT). For 32 hours APF wings, pupae were decapitated and fixed in 4% paraformaldehyde in PBS for 12-18h. All samples were washed in PBS three times and, subsequently, pupal wings were dissected in PBS and immediately blocked for 30min in 1% normal goat serum in PBT. After fixation, all samples were washed three times in PBT, blocked for 30 minutes in 1% normal goat serum in PBT, and incubated overnight at 4C with primary antibodies in PBT. After primary incubation, samples were washed three times in PBT, incubated with secondary antibodies in PBT for 2-16h at room temperature, washed three times in PBT, and mounted in n-propyl gallate mounting medium. Samples were imaged on Zeiss LSM 800 and 880 confocal microscopes. GFP was imaged using the endogenous fluorescent signal. Antibodies: mouse anti-NECD (1:100, Developmental Studies Hybridoma Bank (DHSB)), mouse anti-Delta (1:100, DHSB), mouse anti-Cut (1:100, DHSB), guinea pig anti-Eps15 (1:1000, provided by Hugo Bellen), guinea pig anti-Hrs (1:1000, Hugo Bellen), rabbit anti-Myc (1:1000, Rick Fehon), goat anti-mouse cy3 (1:2000, Jackson Immunoresearch), goat anti-guinea pig cy5 (1:2000, Jackson Immunoresearch), goat anti-rabbit cy5 (1:2000, Jackson Immunoresearch).

### S2 cell staining

S2 cells were transfected with pMT-Notch (150 ng), pMT-Notch^Y2328F^ (150 ng), pMT-Su(dx)-Flag (150 ng), PMT-eYFP-Rab7 (250 ng) using dimethyldioctadecylammonium bromide (DDAB). Cells were subsequently incubated at 25C for 24h before induction. Expression of the PMT constructs was induced with 0.7 mM CuSO4 for 24 hours. For imatinib mesylate treatment, imatinib mesylate solution in water was applied to a culture concentration of 200 μM at the time of induction. For staining, cells were settled on poly-L lysine coated slides for 1 hour, fixed for 15 minutes in PFA+PBT, washed three times in PBT, incubated for one hour at room temperature with mouse anti-NECD antibody (1:100) in PBT+1% normal goat serum, washed three times in PBT, incubated with secondary antibodies in PBT+1% normal goat serum for 2 hours at room temperature, washed three times in PBT, mounted in n-propyl gallate mounting medium, and imaged on a Zeiss LSM 880 confocal microscope.

### Transcription assays

2.25 million S2 cells were transfected with 150 ng of each listed gene construct plus 2 ng of NRE-luciferase reporter, and 40 ng of actin-Renilla control. Blank pMT vector was used to equalize total DNA across transfections. Each biological replicate represents a separate transfection. Cells were lysed in 100mM Potassium Phosphate, 0.5% NP-40, 1 mM DTT, pH7.8 for 1 hour on ice. Lysates were loaded in triplicate into an Autolumat Plus LB 953 luminometer. Firefly and Renilla luciferases were activated in separate reactions using luciferase buffer (10mM Mg Acetate, 100mM Tris Acetate, 1mM EDTA, 4.5mM ATP, and 77uM D-luciferin, pH 7.8) and renilla buffer (25mM sodium pyrophosphate, 10mM Na Acetate, 15mM EDTA, 500mM Na2SO4 500mM NaCl, 4mM coelenterazine, pH 5.0), respectively. The ratio of Firefly RLU to Renilla RLU was averaged across technical replicates, and activation values were normalized to Notch activity.

### Molecular cloning of pGEX-NICD, pGEX-NICD^Y2328F^, and pMT-Notch^Y2328F^

pMT-NotchY2328F was made by subcloning the XhoI-XbaI NICD fragment from pMT-Notch into pBluescript. The Y2328F point mutation was introduced using Quickchange PCR using the following primers: 5’-AAGCAGCCGCCGAGCTTTGAGGATTGCATCAAG-3’; 5’-CTTGATGCAATCCTCAAAGCTCGGCGGCTGCTT-3’. The mutated XhoI-XbaI fragment was cloned back into pMT-Notch to create pMT-Notch^Y2328F^. Wildtype and Y2328F XhoI-NotI NICD fragments were also subcloned into SalI-NotI digested pGEX-4T-2 plasmid to create the bacterial expression constructs.

### In vitro kinase assay

GST-fusion proteins were purified from BL21 E. coli cells as described in (Rebay and Fehon, 2009). Kinase assays were performed using recombinant murine c-Abl (NEB). The amounts of GST-fusion protein to add to the reaction were calibrated using western blots of the purification products with mouse anti-NICD (1:1000) antibody. The relative intensity of fulllength bands was used to make equal dilutions of the GST fusion products. Approximately 1μg of GST-fusion protein, kinase buffer (50mM Tris-HCl, 10mM MgCl2, 1mM EGTA, 2mM DTT, .01% Brij 35, pH 7.5, 200 μM cold ATP), 1μl gamma-32P ATP and 1 μL NEB Abl were incubated for 30 minutes at 30°C. Samples were run on a 10% polyacrylamide gel and transferred to a PVDF membrane using standard methods. The membranes were exposed on a Storm phosphoimager. Protein immunoblot analysis of the same membranes using mouse anti-NICD antibody (1:1000) of the same membranes was then performed using standard methods.

## Acknowledgements

We thank Martin Baron for Su(dx)-Flag and Rab7-GFP S2 cell expression plasmids, Sarah Bray for NRE-luciferase plasmid, Spyros Artavanis-Tsakonas for *UAS-Flag-Dx* flies, Hugo Bellen for Hrs and Eps15 antibodies, Rick Fehon for Myc antibodies, Lucy Godley for imatinib mesylate, and Hitoshi Matakatsu for advice on dissection of pupal wings. We thank members of the Rebay lab, Rick Fehon, Sally Horne-Badovinac, Chip Ferguson, and Aaron Turkewitz for helpful discussions.

---

**Figure 2S.**
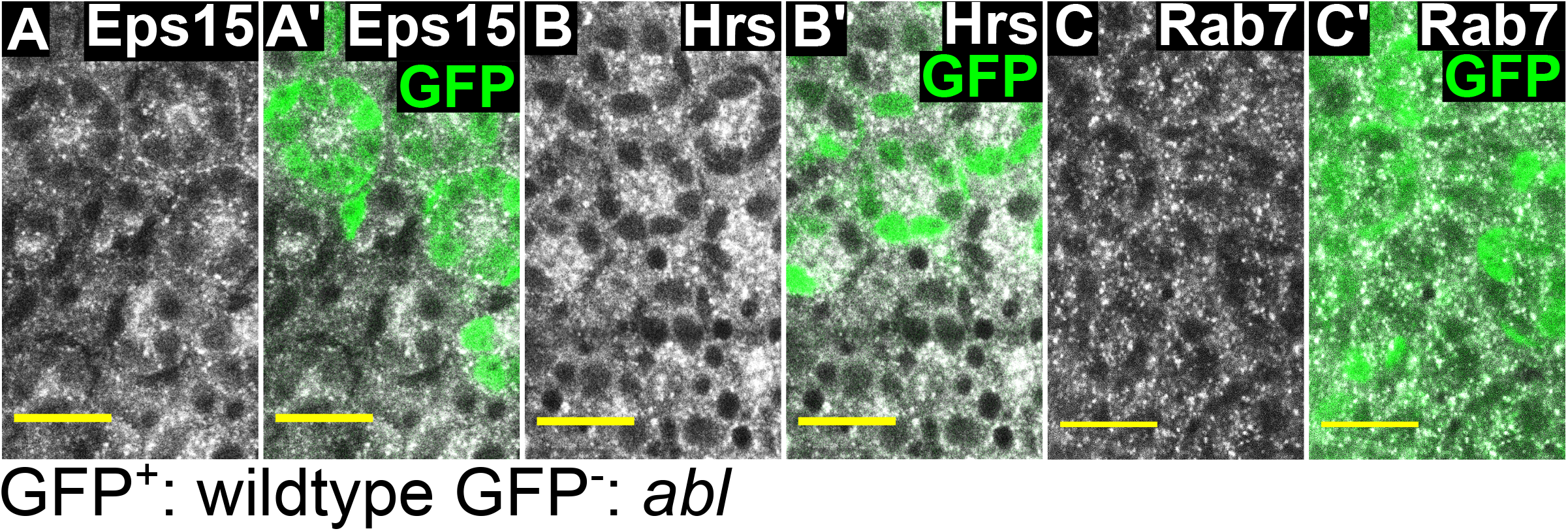
Abl loss does not perturb general endocytic trafficking. (A-C) Panels show single-slice confocal images from 48h APF retinas containing *abl^2^* mutant mitotic clones. Lack of GFP marks *abl* mutant clones in A’-C’. Retinas were stained for the indicated compartment marker. Scale bar = 10 μm. The expression level and subcellular patterns of Eps15 (A, A’), Hrs (B, B’), and Rab7 (C, C’) are the same in wildtype tissue and *abl* clones.

**Figure 3S.**
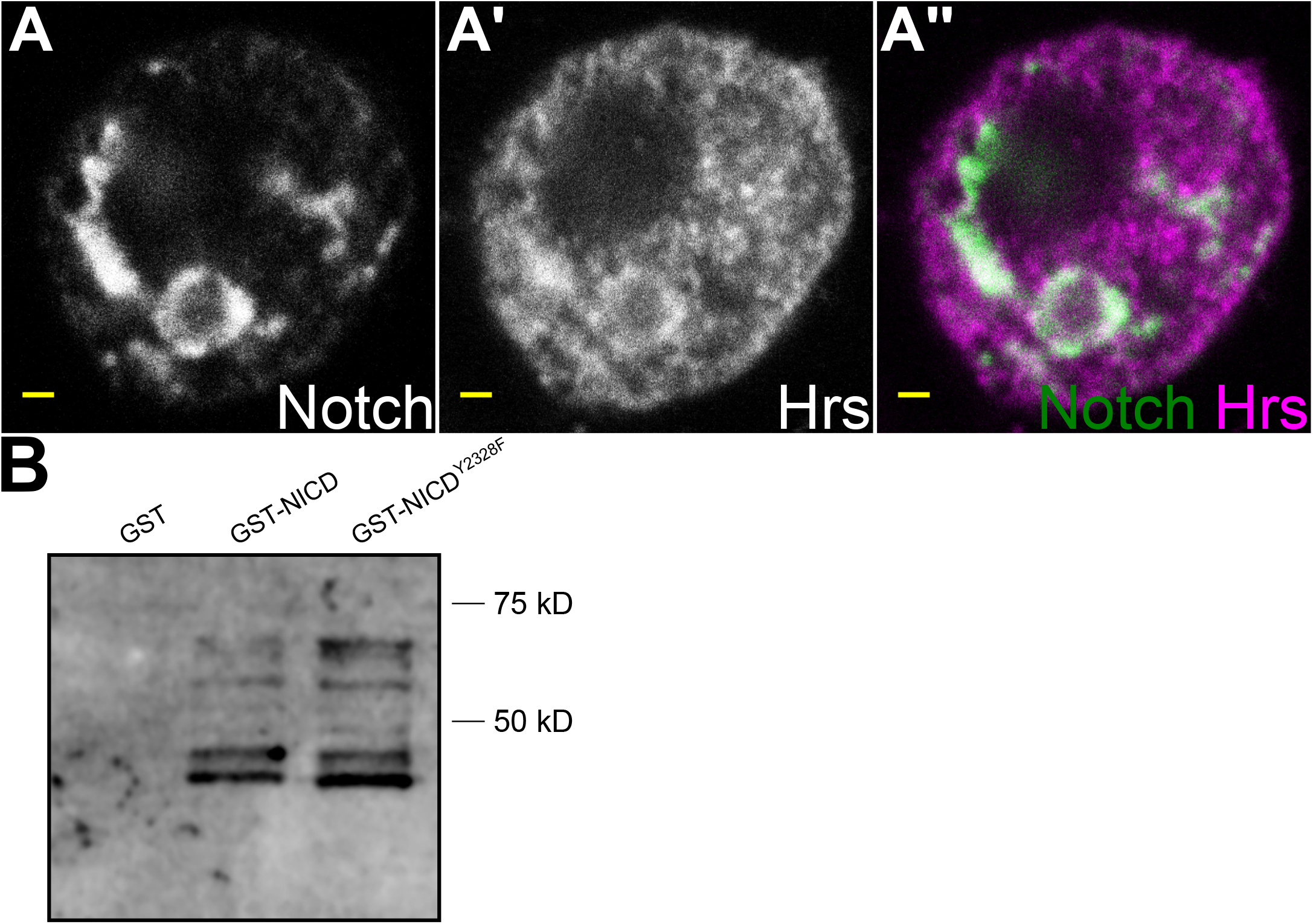
Notch colocalizes with Hrs under imatinib mesylate treatment. (A-A”’) Panels show single-slice confocal images of a representative Notch-expressing S2 cell after treatment with imatinib mesylate for 24 hours. Scale bar = 1μm. Notch colocalizes with Hrs. (B) A western blot of the membrane from Figure 3C using mouse anti-NICD antibody. The antibody specifically recognizes NICD fragments between ~40-70 kD, but does not recognize GST. Note that the amount of full length GST-NICD^Y2328F^ was greater than for GST-NICD, making the reduced phosphorylation of GST-NICD^Y2328F^ shown in Figure 2C even more significant.

